# Estimating the extinction date of the thylacine accounting for unconfirmed sightings

**DOI:** 10.1101/123331

**Authors:** Colin J. Carlson, Alexander L. Bond, Kevin R. Burgio

## Abstract

The thylacine (*Thylacinus cynocephalus*), one of Australia’s most characteristic megafauna, was the largest marsupial carnivore until hunting, and potentially disease, drove them to extinction in 1936. Current knowledge suggests that thylacines became extinct on mainland Australia two millennia prior to their extirpation on Tasmania, but recent “plausible” sightings on the Cape York Peninsula have emerged, leading some to speculate the species may persist, undetected. Here we show that the continued survival of the thylacine is entirely implausible based on most current mathematical theories of extinction. We present a dataset including physical evidence, expert-validated sightings, and unconfirmed sightings leading up to the present day, and use a Bayesian framework that takes all three types of data into account by modelling them as independent processes, to evaluate the likelihood of the thylacine’s persistence. Although the last captive thylacine died in 1936, our model suggests the most likely extinction date would be 1940, or at the latest the 1950s. We validated this result by using other extinction estimator methods, all of which confirmed that the thylacine’s extinction likely fell between 1936 and 1943; even the most optimistic scenario suggests the species did not persist beyond 1956. The search for the thylacine, much like similar efforts to “rediscover” other recently extinct charismatic species, is likely to be fruitless, especially given that persistence on Tasmania would have been no guarantee the species could reappear in regions that had been unoccupied for millennia. The search for the thylacine may become a rallying point for conservation and wildlife biology, and could indirectly help fund and support critical research in understudied areas like Cape York. However, our results suggest that attempts to rediscover the thylacine will likely be unsuccessful.

## INTRODUCTION

The history of conservation biology has included a few exceptional errors, in which experts have pronounced a species extinct only to be “rediscovered.” Perhaps most famous are “Lazarus” taxa known originally from the fossil record (e.g., the coelacanth, *Latimeria* sp., or dawn redwood, *Metasequoia* sp.), but even recently declared species extinctions can also sometimes be overturned; just this year, the “rarest dog in the world,” the New Guinea highland wild dog (*Canis lupus dingo*), was rediscovered after an absence beginning in 1976. Hope of rediscovering an “extinct” species can inspire volumes of peer-reviewed research, and sometimes a single controversial sighting (Fitzpatrick et al. 2005) can be enough to reignite controversy and justify seemingly-endless field investigation, as in the ongoing search for the Ivory-Billed Woodpecker (*Campephilus principalis*) despite all odds (National Audubon Society 2016). Similarly, in Queensland, Australia, two recent unconfirmed sightings have inspired a new search for the thylacine (*Thylacinus cynocephalus*).

The thylacine, or Tasmanian tiger, has been presumed extinct since the last captive specimen died on September 7, 1936 (Sleightholme & Campbell 2016). Thylacines are believed to have gone extinct on the Australian mainland two millennia ago, persisting as Tasmanian endemics (Paddle 2002). State-sponsored eradication in Tasmania between 1886 and 1909 caused a devastating population crash (Sleightholme & Campbell 2016). This eradication campaign, combined with prey declines, could have been sufficient extinction pressure (Prowse et al. 2013), but other research strongly suggests a disease similar to canine distemper could have helped drive the species to extinction (De Castro & Bolker 2005; Paddle 2012). While the mechanism has been a topic of debate, the extinction status of the thylacine has been essentially unchallenged in peer-reviewed literature. Despite this, sightings have continued throughout Tasmania and mainland Australia, often gathering international media attention. In January 2017, two unconfirmed “detailed and plausible” sightings in the Cape York Peninsula of Queensland have sparked renewed interest in the thylacine’s persistence, particularly in the Australian mainland; researchers currently intend to investigate those sightings using camera traps later this year (James Cook University 2017.).

Is there empirical support for this most recent search? Extinction date (τ_*E*_) estimators have been a key part of parallel debates about the Ivory-billed Woodpecker (*Campephilus principalis*); what little work has been done on the thylacine places τ_*E*_ in 1933-1935, with only one model (using temporally-subsetted data) suggesting the species might be extant (Fisher & Blomberg 2012), and a subsequent study revising that estimate to suggest a rediscovery probability of zero after 1983 (Lee et al. 2017b). These methods are sensitive to inaccurate data and false sightings, but recently developed Bayesian models differentiate between the processes of accurate and false sightings explicitly, allowing researchers to include uncertain sightings in models as a separate class of data (Solow & Beet 2014). Here, we apply those models (and several other extinction date estimators) to thylacine sightings, and ask: what is the probability that the species might be rediscovered?

## METHODS

Most of the sightings in our dataset are from Sleightholme & Campbell’s appendix (covering 1937-1980) (Sleightholme & Campbell 2016), which includes 1167 post-1900 sightings classified as a capture, kill, or sighting; and Smith’s (1981) summary of 243 sightings from 1936-1980. Additional sightings were compiled from Heberle (2004), as well as records detailed on public websites maintained by interested citizen groups (www.tasmanian-tiger.com, www.thylacineresearchunit.org, and www.thylacineawarenessgroup.com) supplemented by online news stories from 2007-2016. For each year from 1900-1939, we used the sighting of the highest evidentiary quality, with captures or killed individuals being confirmed specimens (n = 101; **Table S1**). This reduction to at most one record per year was only required by some models, such as that of Lee *et al.* (2014) below, and we aimed for data consistency across methods. Records were scored as confirmed specimens (e.g., from bounty records, museum specimens, or confirmed captures), confirmed sightings (sightings agreed as valid by experts), and unconfirmed sightings (sightings not considered valid by experts; **Figure 1**). The final dataset we assembled spans 1900 to 2016, with the last confirmed specimen collected in 1937; a total of 36 of those years included confirmed sightings. Only one of those years (1932) was recorded with an expert sighting without physical evidence. Because there are also likely unreported unconfirmed sightings, we also ran models assuming that an unconfirmed sighting occurred in every year from 1940-2016, which produces a marginally higher chance of persistence, but without changing the overall conclusions (**Figure S2**).

**Figure 1.**
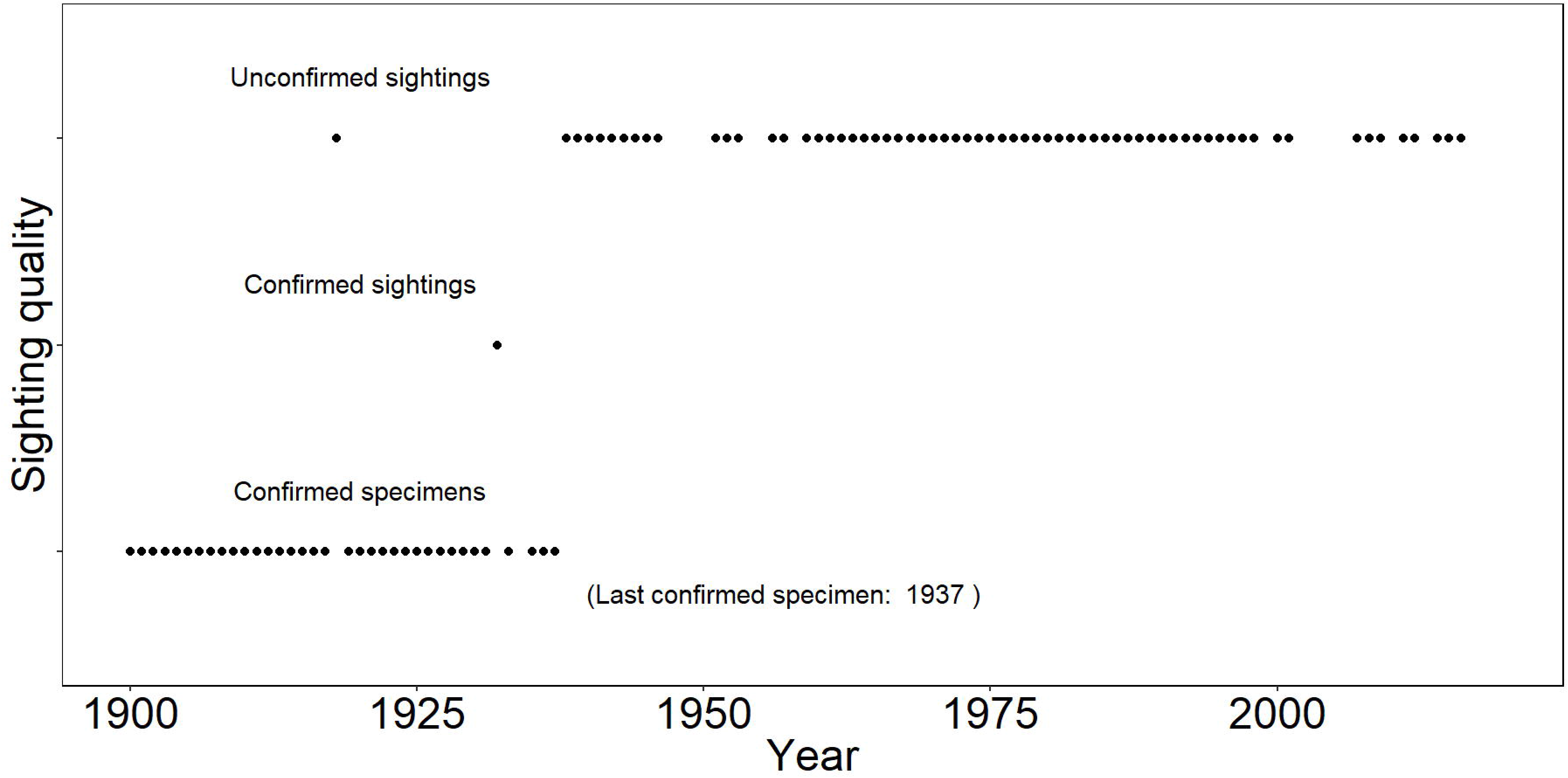
Thylacine sighting data. Specimens are treated as an absolute, certain form of evidence, while expert-verified sightings are treated as an intermediate level of certainty. Controversial sightings, or indirect evidence based on scat or tracks, are classified as unconfirmed sightings, the weakest source of evidence.

For all analyses, we considered the species across its historical range (i.e., mainland Australia and Tasmania), including valid sightings from Tasmania alongside highly questionable sightings from mainland Australia, despite the species’ extirpation two millennia earlier on the continent; we consider this the only optimistic modeling scenario for the thylacine’s persistence in which recent high-profile sightings could be valid, even if it correspondingly represents one of the most biologically implausible scenarios. In the Supplementary Information, we present an analysis using only confirmed sightings from Tasmania, which could be considered a more realistic analysis of the probability the thylacine persisted on Tasmania alone (though this would fail to explain the most controversial recent sightings throughout, primarily, mainland Australia).

### Bayesian Extinction Estimators

Methods to identify likely extinction dates from time series data have been popular in conservation biology since the 1990s, but the majority fail to account for the variability in quality and certainty within most sighting records (Boakes et al. 2015). However, several Bayesian methods have been developed recently that incorporate variable sighting quality and invalid sightings into extinction date estimation. These methods rely on the assumption that the species is extinct (E) given a time series of sightings *t* is given by

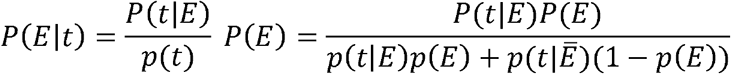

where 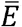 is the scenario that the species is not extinct within the timeframe in question. The prior probability of extinction P(*E*) can be hard to define, though is sometime uninformatively set to 0.5 for explicit estimation of *P(E|t)*. However, the Bayes factor can be used as a test of support for *E* where

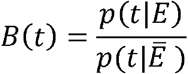

(though, it can be formatted in reverse in some studies, as in the case of Lee *et al.* [2014]). The relationship between the Bayes factor and the probability of the species’ extinction is given as

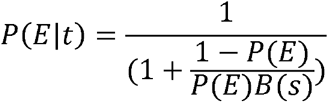

Consequently, with an uninformative prior,

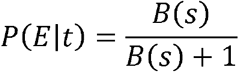

and, with a sufficiently large Bayes factor, the probability of persistence is given by

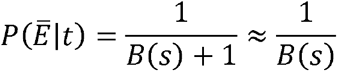

A handful of sighting date methods have been developed using Bayesian frameworks that allow estimation of the Bayes factor and thereby support hypothesis testing. The set of models on which we focus were first developed by Solow *et al.* (2012), who proposed a method in which all sightings leading up to a date *t_L_* (the date of the last certain sighting) were certain and all after were uncertain. In that model, valid and invalid sightings are generated by stationary Poisson processes with different rates, but certain and uncertain sightings had the same rate (Solow et al. 2012). A more recent revision (Solow & Beet 2014) proposed two major modifications. In their first modification (“Model 1”), the same assumptions are made as in the original method, except that uncertain sightings are permitted before *t_L_*. Their second modification (“Model 2,” which we use here) differs more notably, in that it also treats certain and uncertain sightings as independent Poisson processes. This model is especially recommended for cases in which certain and uncertain sightings “differ qualitatively,” as in this study; for example, we note that blurry photographic or video evidence, or crowdsourced sighting records from citizen groups, are a unique issue for later, uncertain thylacine sightings. Therefore, this model is more appropriate than Model 1 (or their 2012 model) for our study.

In Solow & Beet’s (2014) Model 2, while the rate of valid sightings is likely to change leading up to an extinction event, after extinction that rate remains constant (at zero) and all sightings are presumed inaccurate. The sighting dataset *t* occurs over an interval [0,*T*), where 0 ≤ τ_*E*_ < *T*. During the interval [0, τ_*E*_), valid sightings occur at rate Λ while invalid sightings occur at rate Θ, meaning that valid sightings occur at proportion

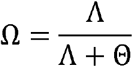

Certain sightings occur, at an independently determined rate, which divides the dataset of sightings *t* into certain sightings *t_c_* and uncertain sightings *t_u_*. The likelihood of the data conditional on τ_*E*_ is given as

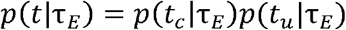

These two values are calculated using *n_c_* (the number of certain sightings, all before τ_*E*_), and *n_u_* (the number of uncertain sightings), where *n_u_*(τ_*E*_) are the subset recorded before τ_*E*_, and ω acts as a dummy variable replacing Ω:

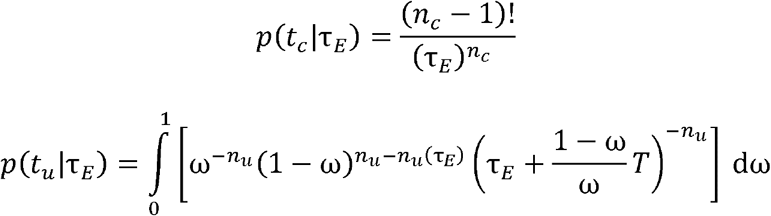

Likelihood *p*(*t*||τ_*E*_) is calculated as the product of those two terms.

The posterior probability that the species became extinct in the interval [0,*T*), which we denote as an event *E* (with alternate hypothesis 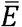), is given for a prior *p*(τ_*E*_) as:

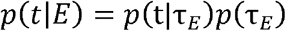

The alternate probability 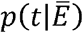 can be calculated by evaluating the same expression given above for *p*(t|τ_*E*_) at τ_*E*_ = *T*. The Bayes factor is given as

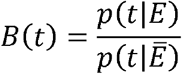

and expresses the relative support for the hypothesis that extinction happened in the interval [0,*T*). The most subjectivity in the method is therefore introduced in selecting the prior τ_*E*_. Solow & Beet (2014) suggest three possibilities: a (1) linear or (2) exponential decline after the last confirmed sighting, or a (3) uniform (uninformative) prior. We elected to use the uniform prior in all the models, as it makes the least constrained assumption about the species’ likely extinction status.

In addition to the models developed by Solow & Beet (2014), we also include another Bayesian model (Lee et al. 2014) that builds on similar foundational work (Solow 1993; Solow et al. 2012). Like Solow and Beet’s (2014) Model 2, the model from Lee et al. (2014) treats certain and uncertain sightings as separate processes, and Lee *et al.* (2014) make slightly different recommendations regarding how to select a prior probability that a given sighting is valid, but the overall intention of the model is largely the same. The approach in Lee *et al.* (2014) is also implemented stochastically using an MCMC approach in BUGS, whereas Solow & Beet’s (2014) model explicitly calculates likelihoods. Here we implement it with some of the simplest possible assumptions: the false positive rate for certain sightings is zero while the false positive rate for uncertain sightings samples from a large uniform distribution. That method can also be more flexibly implemented by assigning different priors to different categories of evidence, as Lee *et al.* (2014) suggest; but rather than use somewhat arbitrarily chosen priors to differentiate among our uncertain reports (a refinement with limited benefits, per a recent study [Lee et al. 2017a]), we simply divide our data into certain and uncertain sightings. A small handful of other Bayesian models exist, but we have included the two appropriate recently-developed Bayesian methods with available code (Boakes et al. 2015).

### Other Extinction Estimators

For the sake of completeness, we also include several other popular and widely-used non-Bayesian estimators with varying levels of complexity (Rivadeneira et al. 2009; Boakes et al. 2015), readily calculated using the R package ‘sExtinct’ (Clements 2013a). Were we to include every unconfirmed, controversial sighting up to 2016, and thereby treat these as valid sightings (as these models make no such distinction), all methods would indicate that the species is unequivocally extant. Consequently, we limit the implementation of other methods to two practical applications, examining how results change by either including (a) only confirmed, uncontroversial specimens and (b) both confirmed specimens and confirmed sightings (**Figure S1**).

Among the methods we include, Robson and Whitlock (1964) suggested a nonparametric method based on the last two sightings:

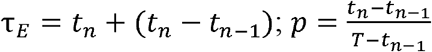

This produced the latest thylacine τ_E_ (see **Figure S1**), as expected, given that the method can be prone to severe overestimation. Burgman *et al.*’s method (1995) uses the length of the period of observation, the number of years with and without records, and the length of the longest consecutive set of years with records to derive a combinatoric probability of unobserved presence. Similarly, Strauss and Sadler (1989) developed a Bayesian method focused on the discrepancy between the observed interval of sightings (between the first and last sighting) and the “true” range of a species in the fossil record. Setting a precedent on which more current methods are based, Solow’s original method (Solow 1993) assumes that sightings are a stationary Poisson process, with the probability of persistence

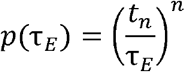

However, a subsequent formulation makes the more accurate assumption that sightings actually follow a truncated exponential distribution, declining until extinction (Solow 2005). Finally, the optimal linear estimator (OLE) method is the most robust non-parametric extinction estimator (Roberts & Solow 2003); based on a subset of the last *s* sightings of *k* total,

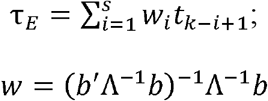

where *b* is a vector of *s* 1’s, and Λ is a square matrix of dimension *s* with typical element

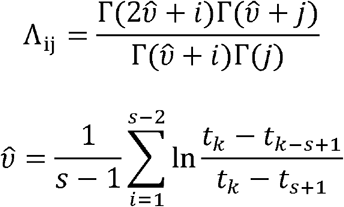

The results of these various analyses are presented in **Table 1** and **Figure S1**.

**Table 1.** Main estimates for thylacine extinction dates. Parametric estimates except the OLE are calculated with a cutoff of α < 0.05; Solow & Beet’s estimate is given by the year with the highest posterior likelihood; Lee’s is given by the posterior probability reaching zero.

| **Model** | **Includes Uncertainty?** | **$\tau_E$** |
| --- | --- | --- |
| Roberts & Solow (2003) | No | 1938 |
| Solow & Beet (2014) | Yes | 1940 |
| Lee et al. (2014) | Yes | 1940 |
| Strauss & Sadler (1989) | No | 1940 |
| Solow (1993) | No | 1941 |
| Solow (2005) | No | 1942 |
| Burgman et al. (1995) | No | 1945 |
| Robson & Whitlock (1964) | No | 1956 |

## RESULTS

The model that differentiates sightings by certainty explicitly found 1940 as the most likely value of τE (with the posterior likelihood declining rapidly thereafter; **Figure 2**), and an extremely low probability that the thylacine could be extant (Solow & Beet Bayes factor = 6.08912 × 10^13^; or, an odds ratio of 1 in 60.9 trillion; using Lee *et al.*’s (2014) method, the probability of persistence is analytically estimated to be zero by 1940). Non-Bayesian estimators all agreed with these findings. Using only confirmed specimens provided an OLE estimated extinction date of 1938 (95% confidence interval: 1937-1943); adding confirmed sightings did not change the estimated extinction date or confidence interval. Other extinction date estimators concur with these findings; Robson & Whitlock’s method (Robson & Whitlock 1964) gives τ_E_ as 1956, the latest estimate (**Table 1; Figure S1**).

**Figure 2.**
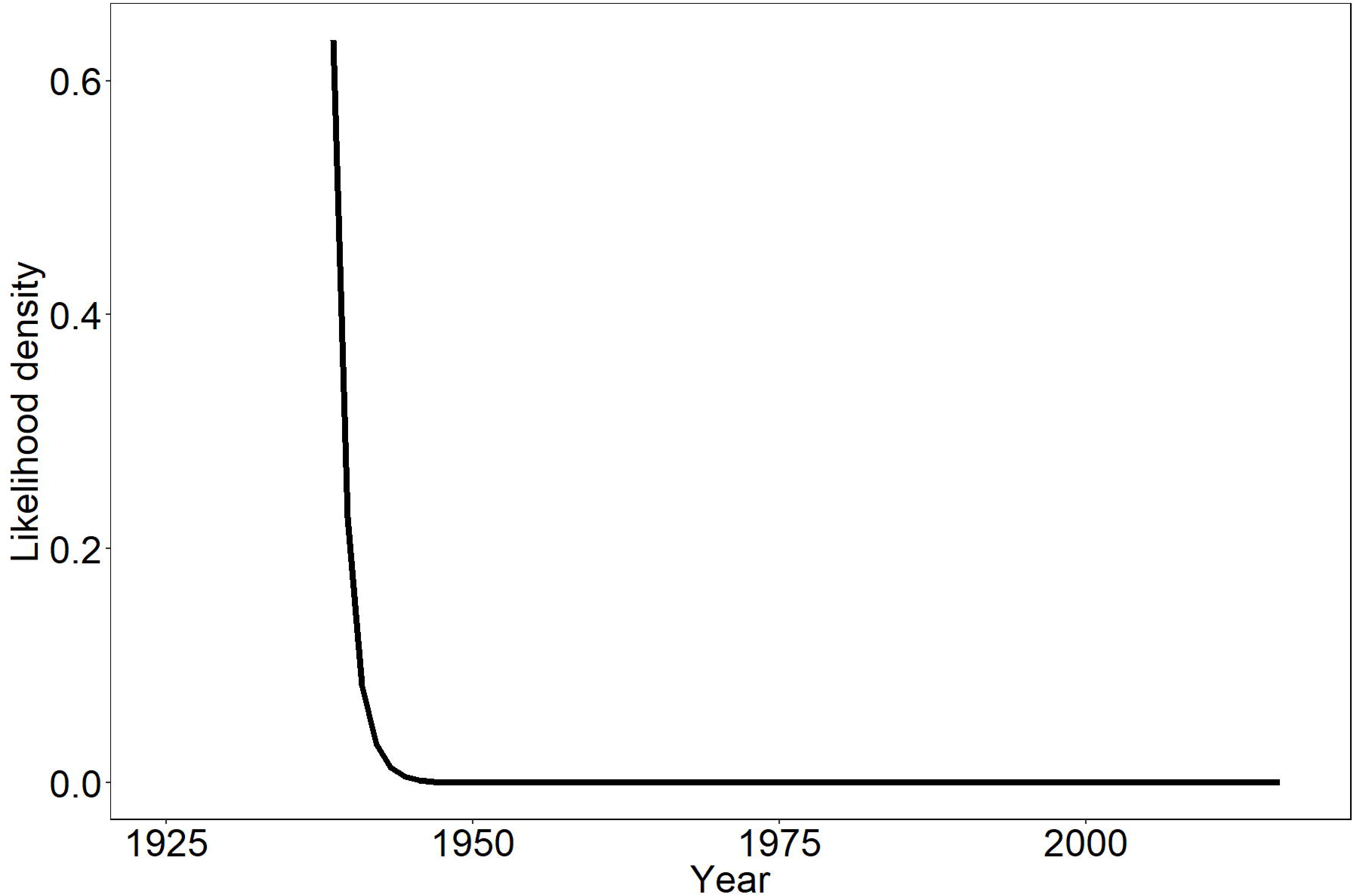
The likelihood of thylacine persistence over time. The figure presents the posterior probability of a given extinction date τ_*E*_ scaled by the area under the entire likelihood curve. In Solow & Beet’s model, specimen-based records are treated separately and as certain observations; consequently, evaluation begins in 1937, the year of the last certain sighting (i.e., extinction prior to that date is not considered).

## DISCUSSION

Based on the results of our primary model, it remains plausible that the thylacine’s extinction could have occurred up to a decade later than believed. But for thylacines to appear in 2017, especially where they are believed to have been absent for two millennia, is highly implausible. The two sightings from Cape York described as “detailed” and “plausible” may be so, from a strictly zoological perspective; but from a modeling standpoint, they fit neatly into a pattern of ongoing, false sightings that follows nearly any high-profile extinction. Models can be wrong, and new data may overturn a century of common knowledge in what could be one of the most surprising re-discoveries in conservation history.

The hope of rediscovering extinct species is one of the most powerful emotional forces in conservation, and can bring attention to threatened species and ecosystems while igniting public interest (and funding) (Clements 2013b). The search for the thylacine may reap those benefits, and the proposed 2017 search has already gathered significant attention from journalists and social media. Moreover, the data that will be collected during the search for the thylacine in Cape York may well be invaluable for other conservation studies. But the ongoing search for extinct species, in the broader sense, likely drains critical funds that the conservation of near-extinct species desperately requires. About 7% of some invertebrate groups may already have gone extinct, at which rate 98% of extinctions would be entirely undetected (Régnier et al. 2015). Globally, 36% of mammal species are threatened with extinction (classified as Vulnerable, Endangered of Critically Endangered), including 27% of native Australian mammals (IUCN 2016), and limited resources can be better spent reversing those declines, than chasing the ghosts of extinction past.

## Acknowledgements

We thank A. Beet for the original Matlab code used in Solow & Beet (2014), A. Butler (Biomathematics and Statistics Scotland) for translating the Matlab code into R, and L. Bartlett for helpful criticism and feedback. We thank two anonymous reviewers for helpful feedback, and particularly for providing useful code adapting the OpenBugs scripts of Lee *et al.* (2014) for our study.

## REFERENCES

Boakes EH, Rout TM, Collen B. 2015. Inferring species extinction: the use of sighting records. Methods in Ecology and Evolution 6:678–687.

Burgman MA, Grimson RC, Ferson S. 1995. Inferring threat from scientific collections. Conservation Biology 9:923–928.

Clements C. 2013a. sExtinct: Calculates the historic date of extinction given a series of sighting events. R package version 1.1.

Clements CF. 2013b. Public interest in the extinction of a species may lead to an increase in donations to a large conservation charity. Biodiversity and Conservation 22:2695–2699.

De Castro F, Bolker B. 2005. Mechanisms of disease-induced extinction. Ecology Letters 8:117–126.

Fisher DO, Blomberg SP. 2012. Inferring extinction of mammals from sighting records, threats, and biological traits. Conservation Biology 26:57–67.

Fitzpatrick JW et al. 2005. Ivory-billed Woodpecker (Campephilus principalis) persists in continental North America. Science 308:1460–1462.

Heberle G. 2004. Reports of alleged thylacine sightings in Western Australia. Conservation Science Western Australia 5:1–5.

IUCN. 2016. The IUCN Red List of Threatened Species. Version 2016-3. Available from http://www.iucnredlist.org (accessed March 25, 2017).

James Cook University. (2017). Press release: FNQ search for the Tasmanian Tiger. Available from https://www.jcu.edu.au/news/releases/2017/march/fnq-search-for-the-tasmanian-tiger.

Lee TE, Bowman C, Roberts DL. 2017a. Are extinction opinions extinct? PeerJ 5:e3663.

Lee TE, Fisher DO, Blomberg SP, Wintle BA. 2017b. Extinct or still out there? Disentangling influences on extinction and rediscovery helps to clarify the fate of species on the edge. Global Change Biology 23:621–634.

Lee TE, McCarthy MA, Wintle BA, Bode M, Roberts DL, Burgman MA. 2014. Inferring extinctions from sighting records of variable reliability. Journal of Applied Ecology 51:251–258.

National Audubon Society. 2016b, April 11. The quest for the Ivory-Billed Woodpecker heads to Cuba. Available from http://www.audubon.org/news/the-quest-ivory-billed-woodpecker-heads-cuba.

Paddle R. 2002. The last Tasmanian tiger: the history and extinction of the thylacine. Cambridge University Press.

Paddle R. 2012. The thylacine’s last straw: epidemic disease in a recent mammalian extinction. Australian Zoologist 36:75–92.

Prowse TA, Johnson CN, Lacy RC, Bradshaw CJ, Pollak JP, Watts MJ, Brook BW. 2013. No need for disease: testing extinction hypotheses for the thylacine using multi-species metamodels. Journal of Animal Ecology 82:355–364.

Régnier C, Achaz G, Lambert A, Cowie RH, Bouchet P, Fontaine B. 2015. Mass extinction in poorly known taxa. Proceedings of the National Academy of Sciences 112:7761–7766.

Rivadeneira MM, Hunt G, Roy K. 2009. The use of sighting records to infer species extinctions: an evaluation of different methods. Ecology 90:1291–1300.

Roberts DL, Solow AR. 2003. Flightless birds: When did the dodo become extinct? Nature 426:245–245.

Robson D, Whitlock J. 1964. Estimation of a truncation point. Biometrika 51:33–39.

Sleightholme SR, Campbell CR. 2016. A retrospective assessment of 20th century thylacine populations. Australian Zoologist 38:102–129.

Smith SJ. 1981. The Tasmanian tiger, 1980: a report on an investigation of the current status of thylacine Thylacinus cynocephalus, funded by the World Wildlife Fund Australia. National Parks and Wildlife Service.

Solow A, Smith W, Burgman M, Rout T, Wintle B, Roberts D. 2012. Uncertain Sightings and the Extinction of the Ivory-Billed Woodpecker. Conservation Biology 26:180–184.

Solow AR. 1993. Inferring extinction from sighting data. Ecology 74:962–964.

Solow AR. 2005. Inferring extinction from a sighting record. Mathematical Biosciences 195:47–55.

Solow AR, Beet AR. 2014. On uncertain sightings and inference about extinction. Conservation Biology 28:1119–1123.

Strauss D, Sadler PM. 1989. Classical confidence intervals and Bayesian probability estimates for ends of local taxon ranges. Mathematical Geology 21:411–427.

